# Bayesian Decoder Models with a Discriminative Observation Process

**DOI:** 10.1101/2020.07.11.198564

**Authors:** Mohammad R. Rezaei, Alex E. Hadjinicolaou, Sydney S. Cash, Uri T. Eden, Ali Yousefi

## Abstract

The Bayesian state-space neural encoder-decoder modeling framework is an established solution to reveal how changes in brain dynamics encode physiological covariates like movement or cognition. Although the framework is increasingly being applied to progress the field of neuroscience, its application to modeling high-dimensional neural data continues to be a challenge. Here, we propose a novel solution that avoids the complexity of encoder models that characterize high-dimensional data as a function of the underlying state processes. We build a discriminative model to estimate state processes as a function of current and previous observations of neural activity. We then develop the filter and parameter estimation solutions for this new class of state-space modeling framework called the “direct decoder” model. We applied the model to decode movement trajectories of a rat in a W-shaped maze from the ensemble spiking activity of place cells and achieve comparable performance to modern decoding solutions, without needing an encoding step in the model development. We further demonstrate how a dynamical auto-encoder can be built using the direct decoder model; where the underlying state process links the high-dimensional neural activity to the behavioral readout. We applied the dynamical auto-encoder model in estimating the intention to verbally communicate of an epileptic participant and their companions. The result shows that the dynamical auto-encoder can optimally estimate the low-dimensional dynamical manifold which represents the relationship between the brain and behavior.

## 1 Introduction

The rapid development of neural recording technologies over the last few decades has enabled the simultaneous recording of neural activity from an ever-increasing number of brain regions. For research groups interested in relating brain activity to higher-level processes, these data are often recorded during some sorts of experimental tasks, together with behavioral or cognitive observations that are influenced by the task [1, 2]. The higher dimension and multi-modality of these data necessitate the development of analytical solutions capable of making statistically robust inferences about the underlying brain dynamics and their relationship to the observed correlates [3, 4]. A wide variety of statistical and machine learning techniques, broadly known as neural encoder-decoder models, have been developed to address this particular type of problem [5-7]. Such models are built in two stages: first, a neural encoder model builds the conditional distribution of observed neural data given the underlying neural correlates (such as movement or cognitive state), and then, newly observed neural data are decoded to estimate those correlates by applying Bayes’ theorem to the encoder model [8-13]. Although traditional neural encoder-decoder models have been successfully applied to gain insights from low-dimensional data [8, 14, 15], they face multiple modeling challenges when applied to high-dimensional data. One of such problems appears in the encoding step, in which the conditional joint distribution of neural data is built. Due to the large dimension of data, it is hard to properly characterize this distribution. Proposed solutions for this distribution are mainly built upon naive assumptions such as the conditional independence of the individual neural data given the correlates. Even with this assumption, it is not always possible to characterize the distributions of neural data and associated noise components (e.g., they may not be stationary), which introduces further complications and difficulties in building the encoder model. The fact that the neural correlates generally have a lower dimension compared to the neural data might help us to address some of the challenges tied to the classical encoder-decoder modeling methodologies [16-19].

In this work, we propose a Bayesian filter solution for the decoder model in which we build the model directly from the neural data ensemble, as opposed to first formulating the encoder model. A specific variation of this modeling approach has been recently proposed by Harrison et al. [20], in which the decoder model is defined as a function of the current-time neural observation. Given this assumption, the Bayes filter solution is derived for the steady-state condition. Here, we introduce a more general framework of this modeling approach, in which the decoder model is defined as a function of current and previous neural data. In our proposed framework, the filter solution accounts for time variability present in the observed data; in addition to that, we derive the maximum likelihood (ML) estimation of the model parameters using a revised expectation-maximization (EM) technique [21].

Recently, new techniques in machine learning like deep neural networks (DNNs) has been used in neuroscience data analysis to address a similar class of decoding and inference problems. DNNs are used to characterize the direct input and output relationship between neural activity and physiological or neurological states [11, 12, 22]. DNNs are flexible models that are able to extract information from high dimensional and complex data. Regardless of the extensive utilization of DNNs in neural encoding/decoding problems, they still have significant pitfalls like lack of generalizability and interpretation [23]. Many techniques are used to address these issues like using dropout [24], or regularization [25]; they even go further by making DNNs’ parameters stochastic by assigning a probability distribution to each parameter of the DNNs, known as Bayesian DNNs (B-DNNs) [26, 27]. B-DNNs maintain their generalizability even when trained by a small number of data and prevent overfitting issues [28]. In this research, we incorporate DNNs and B-DNNs into our framework to add more flexibility and generalizability to the direct decoder model to capture complex dynamics present in neural data.

We apply our modeling framework to decode the 2-D trajectories of a rat moving through a W-shaped maze from the ensemble spiking activity of place cells; our proposed methodology demonstrates decoding results that are comparable to the state-of-art models, a point-process encoder-decoder model [29]. It is worth emphasizing that no encoder model is being built in our proposed modeling framework.

Our proposed Bayesian filter solution can be applied to a broader class of neural encoding-decoding problems, in which the connection between brain dynamics and neural correlates are defined through a low-dimensional dynamical manifold [19, 29-31]. For instance, when behavioral readouts are used as the correlates, the manifold will represent the cognitive states that underlie these behaviors. To address these class of problems, we propose a modeling solution in which a behavioral encoder model is used together with the direct neural decoder model to find the dynamical manifold linking behavior and the underlying neural activity. The proposed encoder-decoder model can be viewed as a dynamical auto-encoder model with the cognitive states as the latent manifold, and the behavior and neural data as different measures of the same dynamical latent structure. We conclude our findings with an application of this solution to a novel decoding problem, in which we seek to decode the communicative intentions of an epileptic study participant (with electrodes implanted for clinical purposes) from their neural activity, by modeling communicative intent as the underlying cognitive state. Whilst we demonstrate our modeling result, we compare our proposed modeling solution with the state-of-art neural decoding solutions including exact point-process [8, 15, 19, 29, 30], generalized linear models [9, 32], and DNNs/B-DNNs [11, 12, 22].

## 2 Problem Formulation

Here, we begin by formulating the direct decoder (D-D) model using a discriminative observation process. We derive the Bayesian filter and parameter estimation solution for this model and expand it to a more generalized form of a *dynamical auto-encoder* model. For the D-D model, we propose a revised EM [21] algorithm that helps us to find a maximum likelihood estimate of the dynamical auto-encoder model-free parameters.

In the state-space modeling solution, care must be taken to build an accurate model of the observation process. We focus on the class of problems for which the dimensionality of neural observations is much larger than the number of state processes, an appropriate constraint for the wide range of problems in neuroscience that deal with multi-electrode neural recordings [33-35].

### 2.A Direct Decoder Model

Let us assume we have *K* observations from *k* = 1 to *k*. Let’s assume ***x***_k_ represents the state (latent process, or underlying cognitive process) at time index *k*, and ***s***_k_ is the observed neural activity (observation process) at the same time index *k*. We define the history term ***h***_k_ as the subset of previous neural observations, ***h***_k_ ⊂ {***s***_1_, …, ***s***_k−1_}. As with a Bayes filter solution, our objective is to estimate

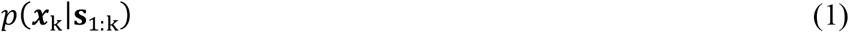

which is the posterior distribution of ***x***_k_ given observed neural activity till time *k*. Using a recursive filter solution [36], the filter update rule at time index *k* is defined by

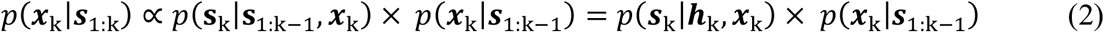

Given the definition of the history term, we can rewrite the filter update rule as

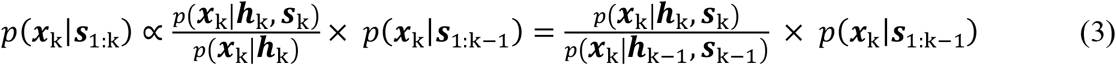

Now, we can build a recursive solution for the update rule using the Chapman-Kolmogorov equation [37]:

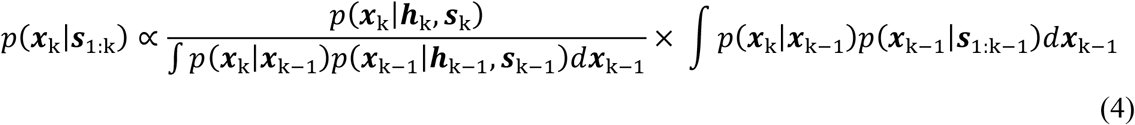

The fraction term on the right-hand-side of equation (4) represents the likelihood function in the standard state-space model. It is the ratio of two likelihood functions for each value of ***x***_k_. The denominator defines the likelihood of ***x***_k_ given the history of observation until time *k* and the numerator is the likelihood of ***x***_k_ when considering the current observation together with the history term. This likelihood can be large or small depending on the information being carried by ***s***_k_ about ***x***_k_, which changes the posterior distribution of ***x***_k_ given the observation until time k. Note that the two Chapman-Kolmogorov equations in equation (4) define the likelihood of ***x***_k_ given two different history terms; these two likelihoods cancel each other if ***h***_k_ = {***s***_1_, …, ***s***_k−1_}. In practice, modeling solutions to characterize *p*(***x***_k_|***h***_k_, ***s***_k_) are misspecified and ***h***_k_ will be limited to a subset of {***s***_1_, …, ***s***_k−1_}. For *p*(***x***_k_|***h***_k_, ***s***_k_) models with a long history term, the two likelihoods become similar, and the filter estimate is being mainly driven by *p*(***x***_k_|***h***_k_, ***s***_k_). When the history term is short, the dynamics of these likelihoods become important in the filter estimation.

As a part of the state-space model, we define the state transition process at time index *k* by

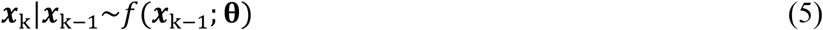

where ***x***_k_ is the cognitive state variable at time index *k* and **θ** is the set of free parameters of the state equation.

Now, we can use the D-D filter solution derived in equation (4) to build the conditional distribution of state for the discriminative model, given the current observation and observation history. We call this the “prediction process”, which is described as

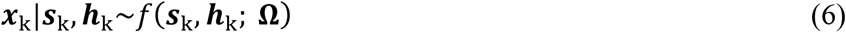

where ***s***_k_ and ***h***_k_ are the neural activity and history term at time index k, and **Ω** is the set of free parameters for the discriminative model.

The D-D model (represented by the schematic in **Figure 1.A**) is comprised of the state and prediction processes defined by equations (5) and (6). With the prediction process, we no longer require an explicit description of the observation process, or the conditional distribution of the observation. The noise process in the prediction process in general is well-defined – note that the noise process for the state is already defined in the state process – and thus the direct decoder model can be easily constructed. The prediction process itself can be modeled using a variety of solutions including a generalized linear model (GLM) [16], a neural network [12], or a linear regression with regularization [38]. It can also incorporate non-linear terms like interaction terms defined by ***s***_k_ and ***h***_k_ along with their higher-order combinations.

**Figure 1.**
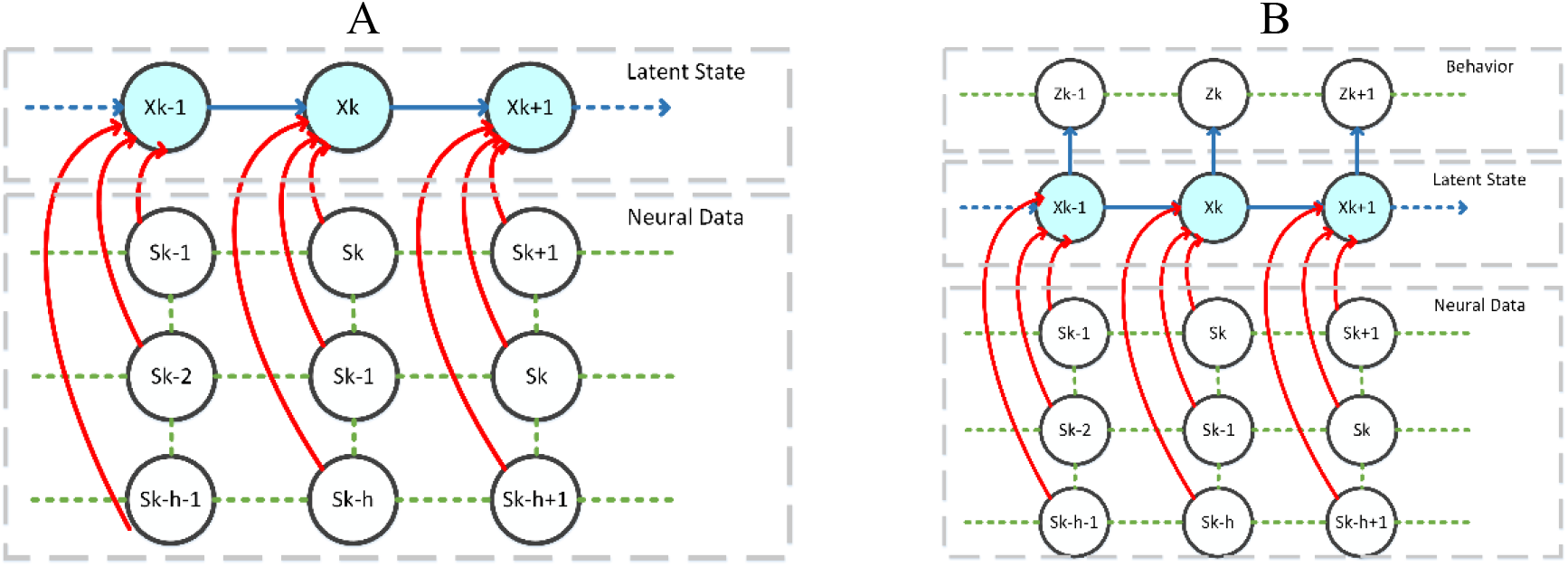
Direct Decoder Model. **A**. Schematic representation of the direct-decoder model. *s*_*k*_ represents neural activity at time index *k, h* represents the number of previous time points in the history term (determined by model selection techniques), and **x**_k_ is the state variable. **B**. Schematic representation of the dynamical auto-decoder model. **z**_***k***_ represents the behavioral readout at time index *k*, which is defined as a function of the state variables. The other parts of the model are the same as the D-D model.

For the model identification step (defined in the next section), we also require a smoother solution of the state which is defined by

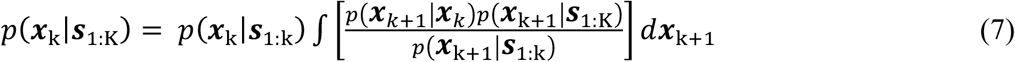

### 2.B Dynamical auto-encoder model

Here, we discuss how the D-D model can be expanded to a dynamical auto-encoder model [19] by incorporating behavioral readouts. Let’s assume that ***z***_*k*_ represents the behavioral observation at time index *k*; given ***z***_*k*_ and **s**_k_, our objective is to estimate

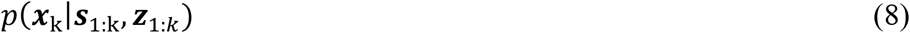

As with the D-D methodology, we can express the filter solution defined in equation (8) as

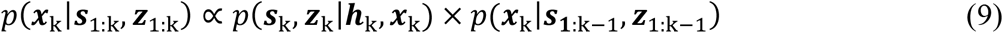

We further assume that the ***s***_k_ and z_k_ are independent given ***x***_k_ [19]; as a result, we can rewrite equation (9) as

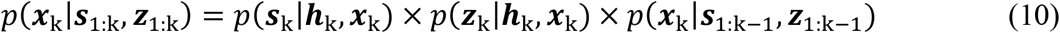

Given the definition of the history term, we can rewrite the filter update rule as

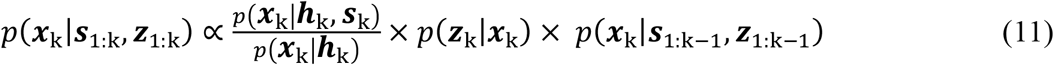

where ***z***_k_, behavioral readout, is assumed to be independent of ***h***_k_ given the state process, ***x***_k_. With this assumption, we can build a recursive solution for the filter update rule [36]:

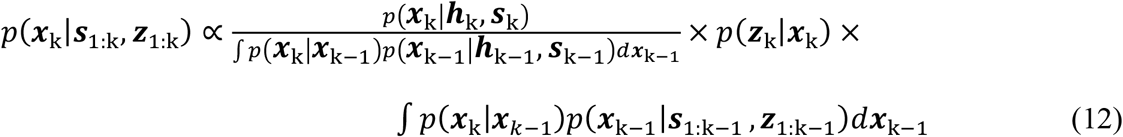

As with the D-D model, we describe the state transition process at time index *k* as a function of the previous state value and **θ**, the set of free parameters of the state equation.

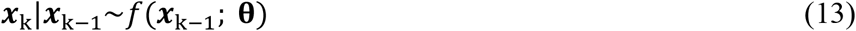

For the auto-encoder model, we have two processes: (1) a prediction process similar to what that has been derived for the D-D model, and (2) a set of observation processes. These processes are described by

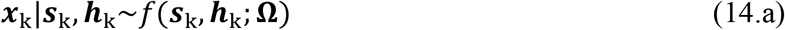

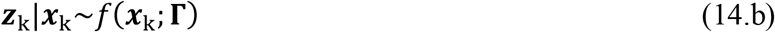

where equation (14.a) is analogous to equation (6) and equation (14.b) is ***z***_k_’s observation process. In (14.b), **Γ** is the set of free parameters that describe the behavioral encoder model. In the D-D and dynamical auto-encoder models described so far, we assume the model parameters and the state dimension are known. In the next section, we describe how the model parameters can be estimated given either ***s***_*k*_ or both ***s***_*k*_ and ***z***_*k*_. In the following sections, we assume the dimension of state process, ***x***_k_, is pre-known. In general, identifying the dimension of the state process is a challenging modeling problem; in the discussion section, we will discuss possible solutions that can be used or developed in the search for an optimal dimension of the state process.

### 2.C Model Parameter Estimation

We use the EM algorithm [39] to find maximum likelihood estimates of the model-free parameters, a subset of {**θ**, **Ω, Γ**}. The EM algorithm is an established solution to perform maximum likelihood estimation of model parameters when there is an unobservable process or missing observations [21]. The other possible solution includes fully Bayesian or Variational Bayes approaches, which can applied to our modeling framework where a Bayesian prior exist per the model parameters [40]. Here, we present the EM solution for the auto-encoder model, given that a D-D model is a specific form of this model. The EM solution recursively estimates the model parameters {***θ***^(r)^, **Ω**^(r)^, **Γ**^(r)^} – here, superscript r is the iteration of the EM procedure, based on an updated posterior distribution of ***x***_0:K_ and parameter estimates from the previous EM iteration, {*θ*^(r−1)^, **Ω**^(r−1)^, **Γ**^(r−1)^}. The EM algorithm includes two steps: expectation (E-step), and maximization (M-step) [41]. The E-step is defined by

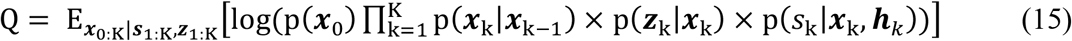

here, the right side is the full likelihood of the state, neural and behavioral readout. The *Q* function can be rewritten as

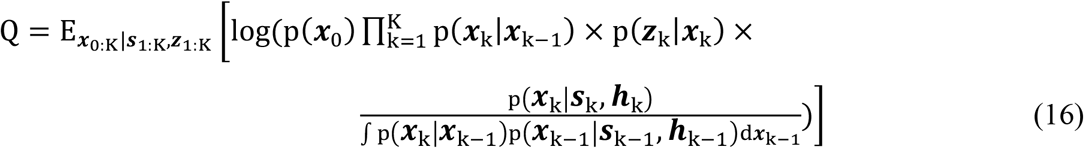

given the prediction process and the behavioral observation process. Expanding the *Q* function yields

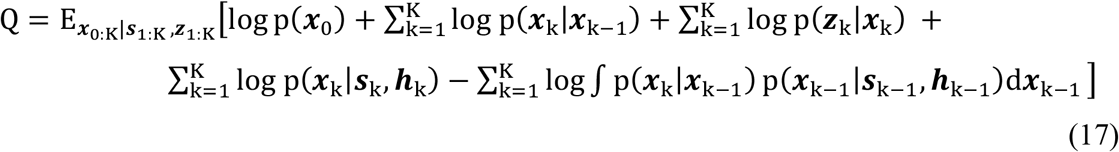

The Chapman-Kolmogorov equation in (17) can be expressed as

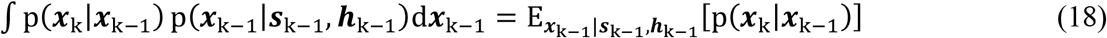

Note that when the state process is linear with an additive Gaussian noise and the prediction process is a multi-variate normal, there is a closed form solution for this expectation. To derive a more general solution, when the state process does not follow a multi-variate normal process, we can use the Jensen inequality – e.g., log(E[g(*x*)]) ≥ E[log(g(*x*))], to exchange the log and expectation operations for the last term in equation (17), which is rewritten in equation (18). This yields a lower bound for *Q*, which can be written as

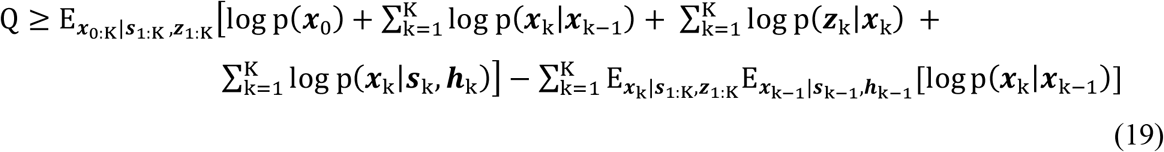

where, the expectation with respect to ***x***_k−1_ in the last term, defined by the prediction process, will be a function of the model free parameters. In the last term of equation (19), the order of log and expectation has changed, and this will make the estimation of Q easier; note that the expectation of log P(***x***_k_|***x***_k−1_) appears twice in the Q function and its expectation is easy to calculate when the state process is a linear multivariate normal.

The updated parameter set at iteration (*r*) can be found by maximizing Q, which is the M-step. This can be done analytically if there is a closed-form solution for the expectation defined in equation (18), or maximizing its lower bound defined in equation (19). The maximization is defined by

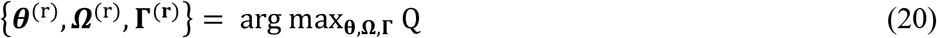

The optimization step in equation (20) can be calculated analytically or numerically, e.g. gradient descent [42]. After each iteration, a new set of parameters are estimated, and the EM routine is stopped when a stopping criterion based on the likelihood of growth or parameter changes is satisfied [21].

### 2.D DNN as direct-decoder model

While a Gaussian linear process can be a proper choice for the direct-decoder model, a more flexible model for the prediction process might capture the complex dynamics presented in high-dimensional data better. To address this, we can utilize DNNs to predict ***x***_k_ distribution given the current and previous neural features. This modeling viewpoint is also aligned with recent advances in the field, where machine learning techniques are used to better understand neural data [11, 12, 22].

A DNN model can be defined by,

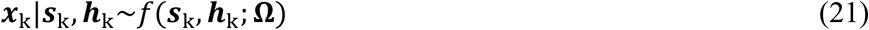

where ***x***_***k***_ is the state process and **Ω** is the network free parameters. This is the same model being used to build the prediction process in equation (14); as a result, we can use DNN in our direct-decoder and auto-encoder model given its free parameters are known. Thus, the objective is to estimate the DNN free parameters in the context of a dynamical auto-encoder model. Note that if ***x***_***k***_ is known, the DNN parameter estimation turns to a supervised problem, which has established solutions [12, 22].

For the auto-encoder model, we applied EM routine to recursively update the model parameters. We can expand this technique to train the DNN by drawing samples from the state posterior distribution. Let’s assume 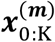 is the *m*^*th*^ sample trajectory from the state given the neural and behavioral data, and we have *M, m* = 1 … *M*, trajectories of the state process. Using the state trajectories, we can calculate *Q* function, defined in equation (19). Using the *M* samples, *Q* is defined by

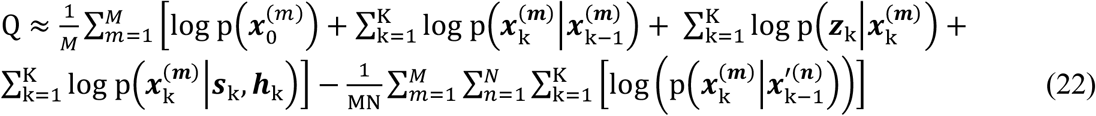

where 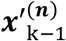 are samples drawn from the prediction process, not the state posterior distribution – note that, we draw *N* samples for it. While Q is maximized, the DNN is trained using samples of the state trajectory, which turns to a supervised learning problem. Note that the DNN is trained on multiple state measure, 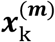, per each time index given ***s***_k_ and ***h***_k_. In other words, the DNN training corresponds to maximize the average of log of likelihood function on multiple samples of state trajectory.

To draw samples from the state trajectory, we can use the conditional distribution of ***x***_***k***_ given ***x***_*k*+1_, ***s***_1:K_, and ***z***_1:*k*_. This distribution can be computed by [43]

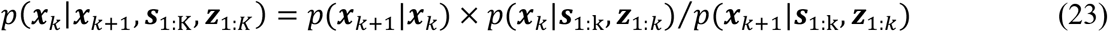

We first draw sample from P(***x***_*k*_|***s***_1:K_, ***z***_1:*k*_), defined in equation (7), and recursively draw samples from equation (23) for time steps *k* − 1 to 0. We then use these samples in equation (22) to find updated model parameters maximizing Q.

In the previous section, we discussed a numerical solution for the filter and smoother steps. In cases where the state is high-dimensional and computing the integrals in filter and smoother equations using simple numerical methods become computationally expensive, we can use sequential Monte Carlo (SMC) methods, otherwise known as particle filters, as an alternate approach for filter and smoother estimation [44, 45]. If we use the particle filter, we can use smoother samples in the Q function defined in equation (22).

Here, we described a DNN with a fixed set of parameters. In order to avoid possible overfitting issues in our DNN training, it is suggested to use a B-DNN in the auto-encoder model [26]. In B-DNN, there will be a probability distribution over the network weights instead of fixed weights. It is possible to build a fully Bayesian auto-encoder model, where not only DNN’s weights are probabilistic but also the observation and state processes parameters are defined through prior distributions. In our modeling solution, we have already derived a fully Bayesian solution for the state process and a maximum likelihood estimate for model-free parameters, including the DNN weights. Extending the DNN parameter estimate to MAP estimate is easy, and it can be done through a penalized likelihood estimate in the context of our EM algorithm [46]. However, solving a fully Bayesian solution for DNN is generally a complex and computationally intractable modeling problem. In Appendix A, we provide a suboptimal solution based on the EM solution we already developed here. The proposed solution is based on the MCMC solution and it might provide a reasonable solution when DNN has a limited number of wights or the network weights are significantly correlated.

## 3 Datasets

We applied our methodology to different decoding problems. In the first problem, the goal is to decode the movement trajectory of a rat from invasive neural recordings, while the rat is foraging in a W-shaped maze for food (Hippocampus dataset) [12, 22, 29, 30]. The second problem investigates how invasive neural recordings and behavioral observations from a human participant can be processed to infer a dynamical internal cognitive process, representing the human participant’s intent to communicate with a companion. In the following section, we describe each dataset in more detail.

### 3.A Movement Dataset

In this dataset, we seek to decode the 2-D movement trajectory of a rat traversing through a W-shaped maze from the ensemble spiking activity of 62 hippocampal place cells [29]. The neural data were recorded from 62 place cells in the CA1 and CA2 regions of the hippocampus brain area of a Long-Evans rat, aged approximately 6 months. The rat has been trained to traverse between the home box and the outer arms to receive a liquid reward (condensed milk) at the reward locations. **Figure 2.A** shows the maze structure and the rat’s movement trajectory in 2-D spaces, where the rat position at each time step is represented by (x, y) coordinates. The spiking activity of these 62 units was detected offline by choosing events that their peak-to-peak amplitudes were above a threshold of 80uV in at least one of the tetrode channels (see **Figure 2.B**). In the experiment process, the actual rat’s position was measured by video tracking software which was used as the ground truth for the position (see **Figure 2.C**). We used a 15-minute-long recording of the experiment, with a time resolution of 33 milliseconds, to analyze different decoding solutions. The first 85% of the recording (∼13 minutes) was used to train the prediction and state processes’ models, and the remaining 15% (∼2 minutes) of the data was used to test the model’s decoding performance.

**Figure 2.**
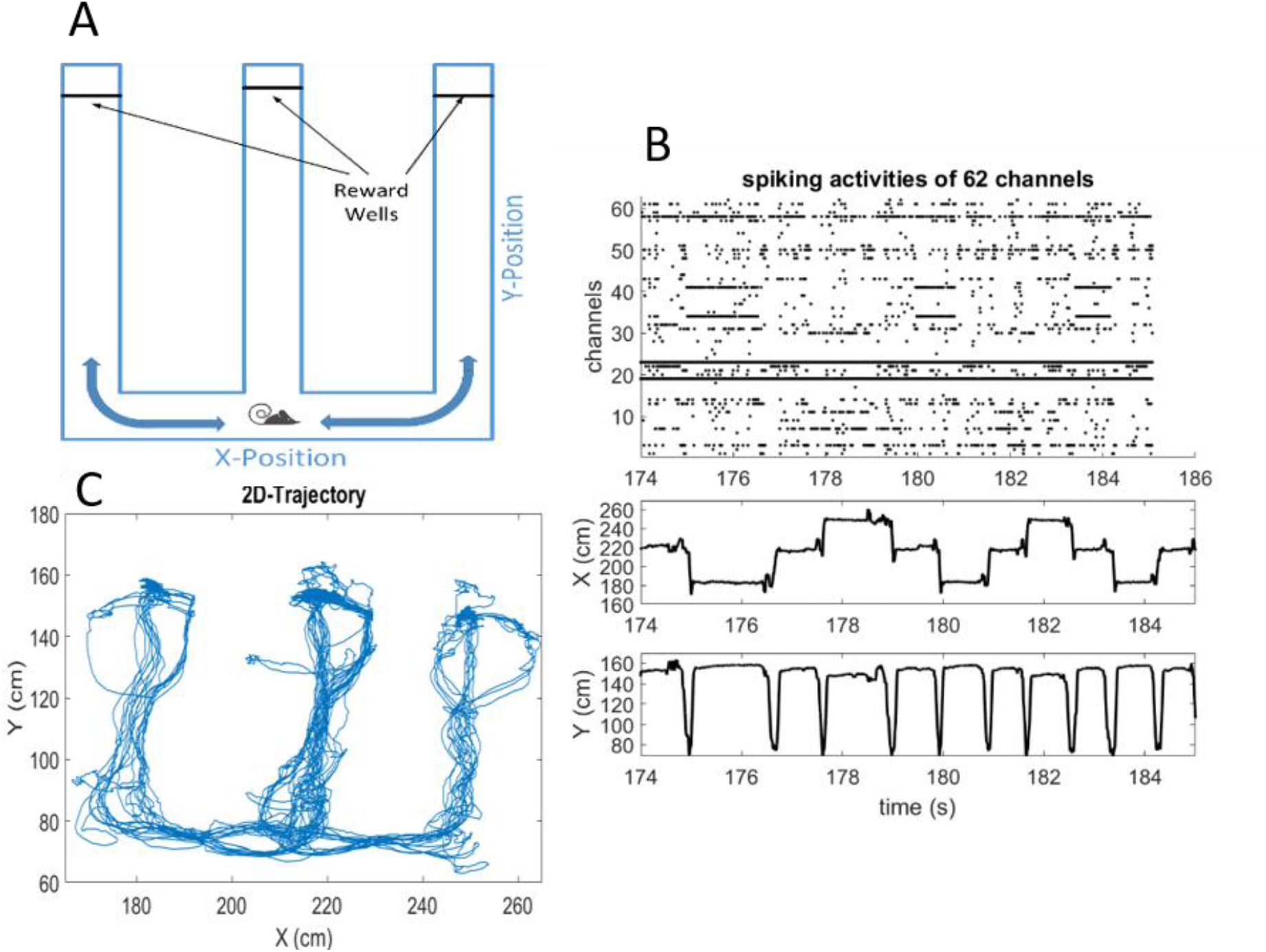
Movement Dataset: the maze topology, the rat movement trajectory, and sample neural data. **A**. W-maze topology, the rat moves from the center arm to the left and right arms to get a food reward. **B**. Both movement trajectories along with *x* and *y* directions and neural activity of the 62 channels. **C**. 2-D trajectory of the rat movement during the experiment. The experiment is about 3 minutes, and during this 3 minutes, the rat has traversed 3 times inside the maze.

### 3.B Conversation Dataset

In this experiment, we investigated how the neural recordings and spoken words from a human study participant can be processed to infer the dynamics of a cognitive state related to verbal communication. In this dataset (unpublished data; Hadjinicolaou, Cash, et al., Massachusetts General Hospital), study participants were implanted with intracranial (sEEG) electrodes for clinical monitoring of their epilepsy, for the duration of their stay in the telemetry ward. The raw neural data were acquired at a sampling rate of 2 kHz using a 128-channel neural signal processor recording system (Cerebus, Blackrock Microsystems, UT) and neighboring channels were re-referenced with a bipolar montage to mitigate volume conduction [47]. The spectral power was estimated for each channel for the theta (4-8 Hz) and gamma (70-115 Hz) frequency bands (Chronux). Neural activity was recorded during conversations between the participants and their companions, including hospital staff, family, friends, and study investigators. All spoken dialog within each recording interval was captured and transcribed to yield individual word timings that are synchronized to the neural data (see **Figure 3**). We used this data – neural activity and spoken dialogs – to examine our dynamical auto-encoder model in search of the underlying intent state of the patient to communicate.

**Figure 3.**
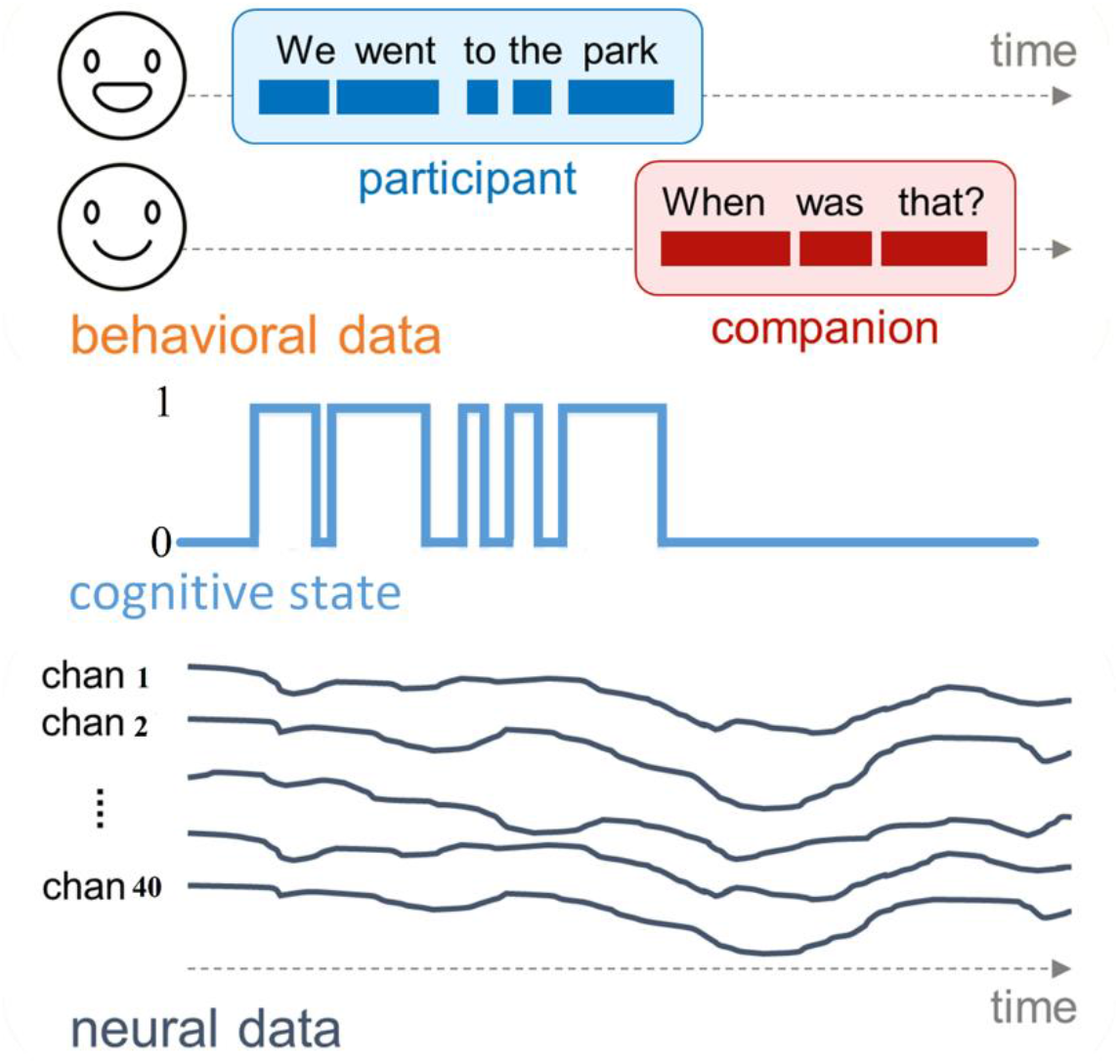
Explanation of the conversation dataset. The diagram shows the relationship between the neural recordings, spoken words from a human study participant, and the cognitive state related to verbal communication during the experiment period.

### 3.C Decoding Problems

Having the direct-decoder modeling framework, we now discuss how the model can be used to decode two different correlates of neural data already described. We describe model identification for these two decoding problems and use two metrics to compare the performance of our proposed solution along with other established decoding techniques. The performance metrics include: mean-squared-error (RMSE), and 95% highest posterior density (HPD) region [48].

#### 3.C.1 Decoding 2-D movement trajectories using the direct-decoder model

We assume the rat’s position in the maze at time interval *k* is specified by the state variable ***X***_k_ = (*x*_k_, *y*_k_), where *x*_k_ and *y*_k_ represent the rat’s *x*- and *y*-coordinates. We define the state process by

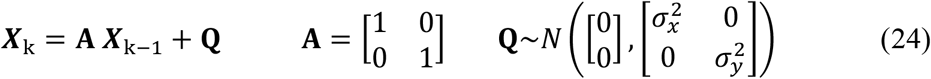

where the covariance matrix **Q** is assumed to be diagonal with 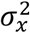 and 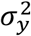 terms representing the movment variances. We estimate these two variances empirically using the rat’s movement during the training session. For the prediction process, we assume the state ***X***_k_ can be predicted using a linear regression model where the predictor variables are the ensemble spiking activity of 62 cells, where each spike train is filtered with a Gaussian window with length 20 ms. We further assume the noise process follows a normal distribution. We build two regression models; one per each coordinate, as a function of ensemble spiking activity. The prediction process and its decomposition into two predictor models are defined by

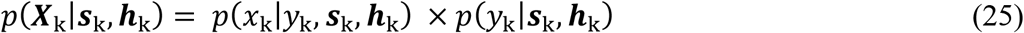

where ***h***_**k**_ represents the history of the ensemble spiking activity from the previous time intervals. Note that given the rat’s movement is bounded by the maze, the state process defined in equation (24) is a misspecified model. To address this issue, we add a penalty term to the prediction process which accounts for the topology of the maze – a detailed explanation of the penalty term can be found in our previous work [29]. The revised prediction process is expressed as

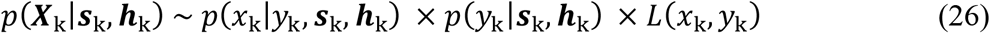

where *L*(*x*_k_, *y*_k_) is close to zero for *x-y* coordinates outside the maze area and one otherwise (**Figure 4.F**). Note that adding the penalty term does not change any aspects of the modeling pipeline.

**Figure 4.**
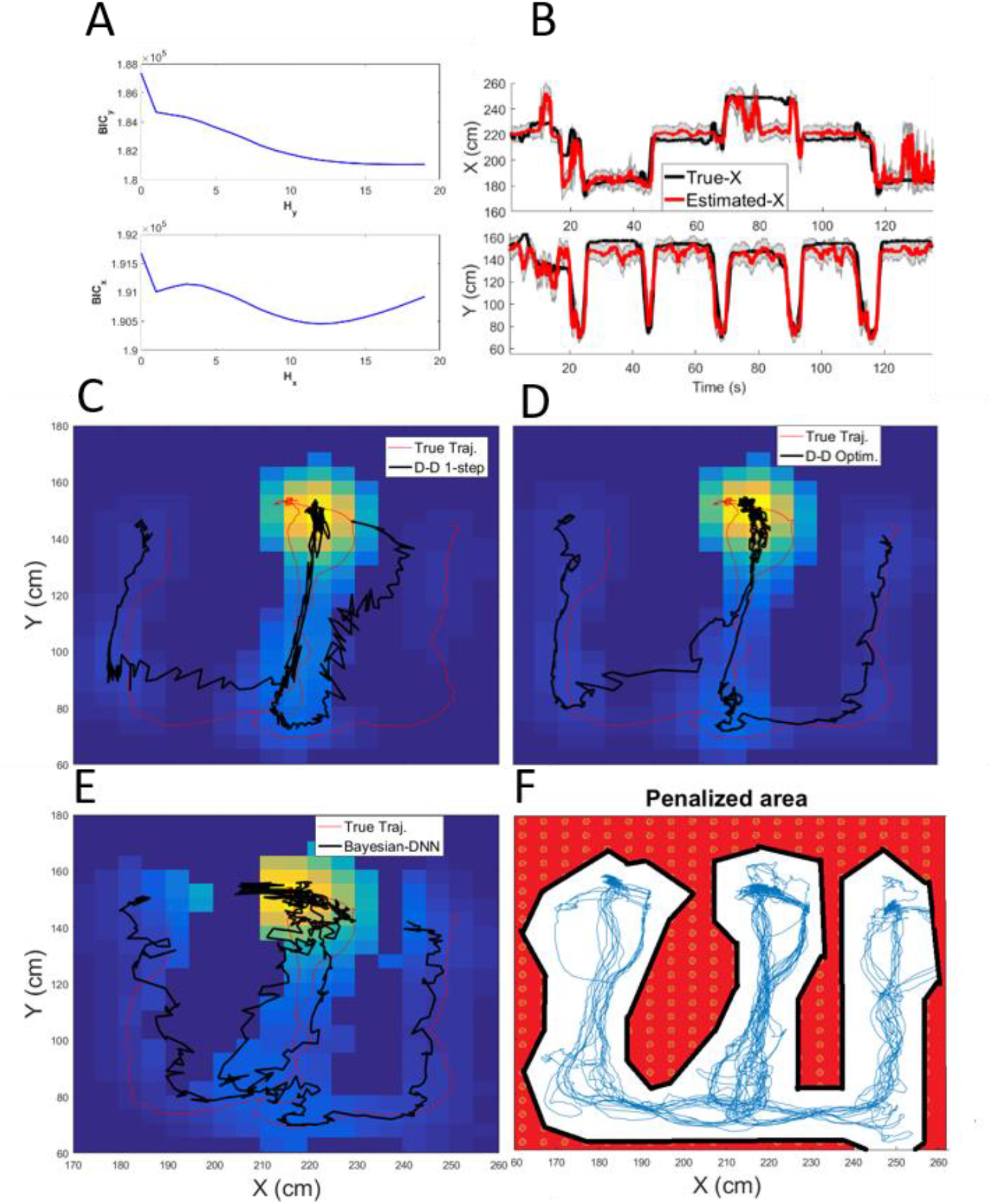
Decoding results in 2-D movement trajectory. **A**. BIC curve for different *x*- and *y*-coordinate model history term lengths. The optimum lengths (corresponding to the minimum BIC) for *hx* and *hy* are 12 and 18, respectively. **B**. The D-D model result for *x* and *y* coordinates with optimal history lengths. **C**. The D-D model result with one-step history terms. **D**. The D-D model result with optimal history term lengths. **E**. The B-DNN model results with optimal history term length. **F**. Visualization of rat movement trajectories (blue traces) together with an overlay of the penalty term *L*(*x*_*k*_, *y*_*k*_) (red area), defined in equation (30). The penalty term is close to zero for *x-y* coordinates outside the maze area (identified by red dots) and one otherwise.

To find the optimal length of the history term for the *x*- and *y*-coordinate regression models, we use a forward model selection process. For the *y*-coordinate model, the null model is defined by the current observation of the spiking activity and the ensemble spiking activity of the previous time points are added recursively. We use a BIC criterion [42] to determine when the increase in the length of the history term does not improve the model fit. For the *x*-coordinate model, the null model is *y*_k_, and the ensemble spiking activity of the current and previous time points are added to the model in search of the optimal length of the history term. As it was done for the y-coordinate, we use a BIC criterion to find the proper length of the history term for the *x*-coordinate model. For this dataset, the history term for the *x*-coordinate ends up to 12 previous time points (∼ 400 milliseconds) and the *y*-coordinate includes 18 time points (∼ 600 milliseconds) (**Figure 4.A**). To assess the performance of our modeling framework, we decoded the rat movement trajectory in the test dataset using four different models: (1) an exact point process decoder model, described in [29], (2) a D-D model with one-step history terms (**Figure 4.C)**, (3) a D-D model with optimal history term lengths (**Figure 4.D** and **Figure 4.B**), and (4) a B-DNN model with optimal history term length (**Figure 4.E**). The performance results in **Table. 1** shows comparable performance between the (exact) point process model and the D-D model with the optimal history term. The performance result also suggests the necessity of incorporating proper history terms in building a more robust decoder model. We expect B-DNN to have a better prediction accuracy compared to other D-D models; however, this is not the case. A possible reason is a limited dataset we have to train the B-DNN model, and thus, the model performance drops in the test dataset despite attaining a superb performance in the training dataset.

**Table 1.** Different Models Decoding Performance.

| Method | RMSE (cm) |  | 95%HPD |  |
| --- | --- | --- | --- | --- |
|  | Train | Test | Train | Test |
| Exact numerical solution | -- | 12.7 | -- | 85.8% |
| D-D result with hx=1 and hy=1 | -- | 17.7 | -- | 85.9% |
| D-D result with optimum hx and hy | -- | 13.8 | -- | 88.7% |
| B-DNN | 14.5 | 15.1 | 88.7% | 88.4% |

#### 3.C.2 Decoding cognitive state using the dynamical autoencoder

The onset times of words spoken by the participant comprise the behavioral signal *z*_*k*_, which is specified as a point-process observation model [15]. The auto-encoder model aims to build the relationship between intention state *x*_*k*_, and neural features, in conjunction with the behavioral observation. The neural features ***s***_*k*_ consist of spectral power at different frequency bands from a subset bipolar recording channels, which add up to 40 features [49].

We assume the prediction process follows a normal distribution, where the expected value is a linear function of the neural features [19]. The intention state *x*_*k*_ is characterized by a random walk model [14]. The auto-encoder model is defined by

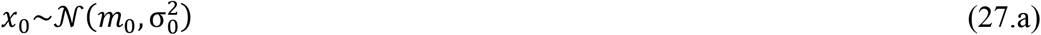

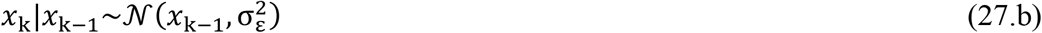

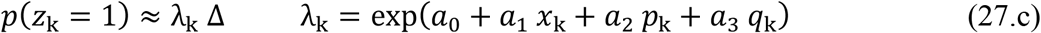

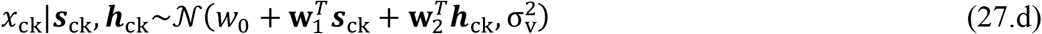

where, 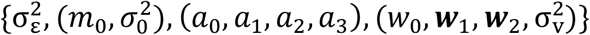 are the model free parameters – to avoid model identifiability issue, we set *a*_1_ to 1. Here, we assume the history term for the conditional intensity of spoken words can be defined by *p*_*k*_ and *q*_*k*_, that corresponds to the number of spoken words from the participant and their companion respectively, over the last 400 milliseconds [8, 50]. The processing interval Δ is set to 50 milliseconds; small enough that the probability of more than one word inside each interval is negligible [15]. Neural features are updated at a slower rate than the behavioral one; they are updated once every 2 seconds. As a result, we have a multi-rate auto-encoder model, and this has been addressed with the *c* term in equation (27.d). The *c* term is equal to 40 – e.g., 2 seconds/50 msec; this implies that we have neural activity updated for every 40 times updates of behavioral data.

We use our modeling solution to estimate the state process, which simultaneously maximizes the likelihoods of the behavioral readout and neural recording. **Figure 5.B** shows the intent state estimation given behavioral readout z_k_ (see **Figure 5.A)**, neural activity ***s***_k_, and optimal history term *h*_k_, which is selected by the LASSO regularization method [38, 49]. The decoding result using only neural activity is shown in **Figure 5.B**. The intention state increases when the participant starts to talk and it decreases when they stop or the other companion starts to talk – or, the participant starts to listen, which is aligned with our expectations. As it can be seen from **Figure 5.B**, by adding the behavioral signals to the decoder model, auto-encoder model, we get more salient state estimation compared to the state estimation run only by the D-D model.

**Figure 5.**
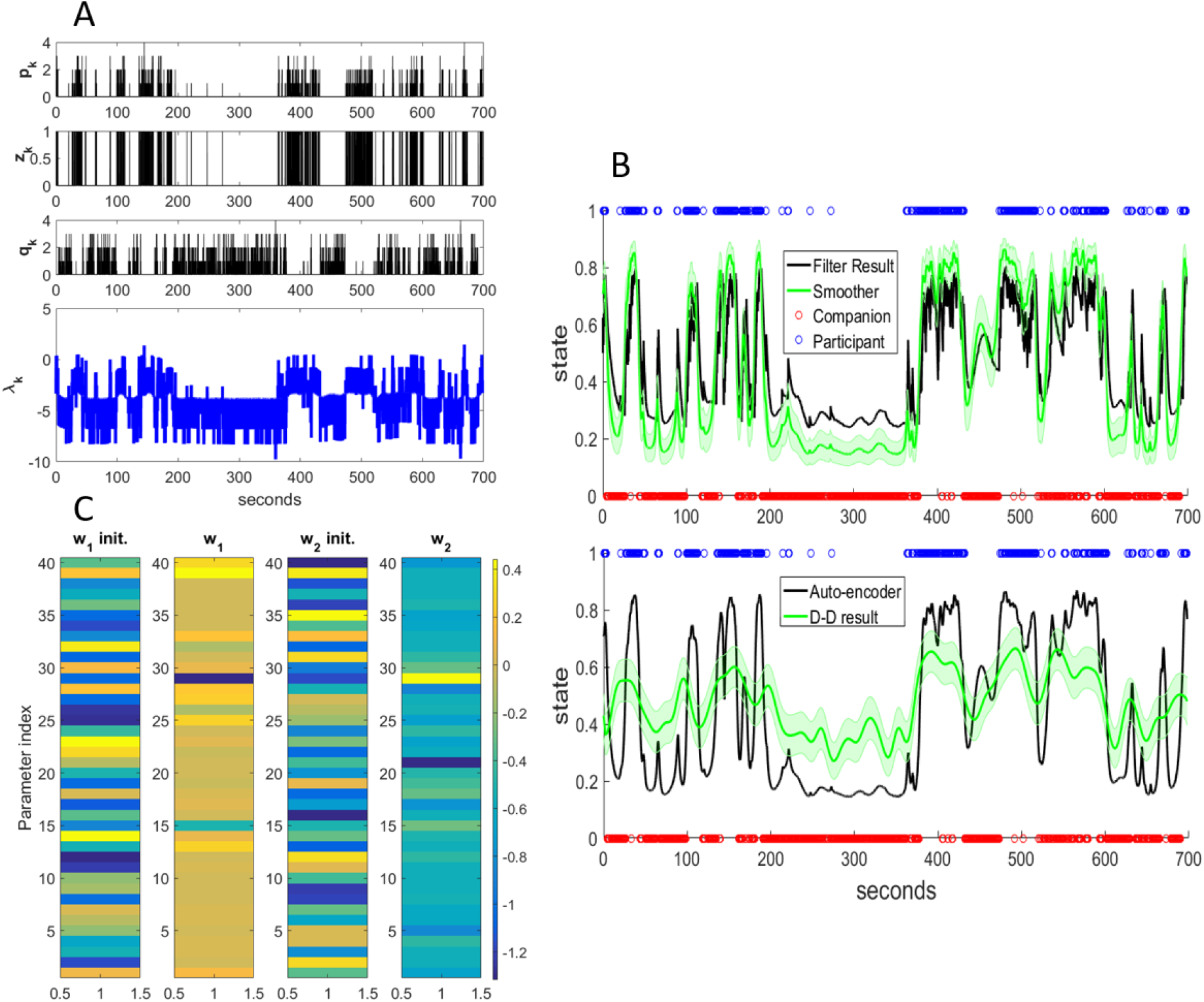
Decoding communication intent using D-D model. **A**. Temporal evolution of *p*_*k*_, *z*_*k*_, *q*_*k*_, and *λ*_*k*_ variables, defined in equation (27). **B**. Conversation intent estimation using the dynamical auto-encoder model. (top) Estimated intent state using both behavioral readout *z*_*k*_, and neural activity *s*_*k*_; (bottom) Estimated intent state using only neural activity. **C**. MLE estimate of the direct-decoder model parameters.

We use the model parameters using the auto-encoder model which is optimized by using both behavioral and neural data. We update the model parameters using the EM technique described in section 2.C. Initial values and optimized values of the D-D model parameter using the EM algorithm are shown in **Figure 5.C**. We assume the history term is 1, and find a sparse set of the weights. This corresponds to a MAP estimate in EM with a Laplace prior to D-D weights. The D-D model weights can also reflect physiological mechanisms of intention, like which neural activity is positively/negatively correlated with the intention and which neural activity is not predictive of the intention – this is not the scope of this research.

## 5 Discussion

In this paper, we introduced a state-space decoder model with a discriminant observation process. The discriminant observation process called D-D model characterizes the state process as a function of current and previous neural data. The model filter solution accounts for time variability present in the observed data; in addition to that, we derive the maximum likelihood (ML) estimation of the model parameters using a revised EM technique [21]. A distinct difference between our work and previous work like those being suggested by Harrison et al. [20] is generalizing of the discriminat process by including the history terms. For this, we not only show the necessity of the history terms through mathematical derivation but also demonstrate the need through a neural decoding problem – e.g., 2-D decoding problem. We then expanded the D-D model to a dynamical auto-encoder model, which lets us link the behavioral readout and high-dimensional neural recording in pursuit of a low-dimensional manifold representing underlying emotional or cognitive states. We discussed how DNNs, including B-DNNs, can be added as the direct-decoder model, which increases the flexibility of the framework to characterize complex dynamics present in the high-dimensional data. Not only we demonstrated how DNNs can be added in the auto-encoder pipeline, but also provided training procedure for DNNs and B-DNNs – Appendix A. Finally, we demonstrated the application of the auto-encoder model in a novel and potentially complex clinical experiment and showed that participant communication intent can be estimated through neural data.

Whilst characterizing the relationship between brain dynamics and behavioral or physical correlates like movement in the context of high dimensional recording is of great importance; a more significant research direction is to estimate underlying cognitive or mental processes that shape the relationship between distributed neural activity and behavior. The dynamical auto-encoder model proposed in this work is well suited for this research objective and can be applied to complex and novel tasks like the communication intent task that we discussed here. The decoding examples presented here highlight the flexibility of our proposed modeling framework, and the fact that, it can be applied to different modalities of neural and behavioral data across different tasks and domains.

In our previous research – Yousefi et al. [19], we discussed two-step encoding and decoding solution to characterize the relationship between brain dynamics and behavior thorough a dynamical cognitive process. In that work, we first estimate the underlying cognitive states using behavioral readout and we then utilize the estimated cognitive state to build a neural encoder and potentially decoder. There, the temporal dynamics of the cognitive state are mandated by the behavior and the neural encoding process looks for neural features that represent dynamics of cognitive state-driven solely by behavior. In dynamical auto-encoder solution; the underlying cognitive state dynamics is not solely driven by the behavior readout; instead, it is defined through joint neural and behavioral readout. This viewpoint can improve our estimation of cognitive state, where different processes come to an agreement in their estimation of the latent dynamical process representing cognitive or emotional states. It is worth mentioning, we already benefit from the D-D process as a part of the dynamical auto-encoder model letting us avoid building many auto-encoder models, each per neural activity.

In both experiments, we used a Gaussian process to build the parametric D-D model. Note that we have a priori knowledge on the observation process noise model, as we define what state process model constructs the underlying dynamical process. The state processes in both decoding problems are defined by Gaussian processes; as a result, we used a Gaussian process for the observation process. The noise process for both decoder problems are assumed to be stationary, whilst the noise process characteristics can change over time. To address this, we can build a more flexible model like DNN to capture changes present in the noise process. Parametric models like those used in two experiments might fail to capture the complex and time-varying dynamics present in the state processes, and using DNNs with their non-linear mapping will boost the model prediction power. Our proposed framework shows how DNNs and B-DNNs can be incorporated into the modeling framework, and how they can be trained as well. A challenge with DNNs is their higher level of flexibility, which will lead to an overfitting problem. To address this, we discussed how to utilized B-DNN in our dynamical auto-encoder model. We also provided the training procedure – Appendix A – to find the posterior distribution of DNN weights, which to the best of authors’ knowledge is novel and can help to build more robust decoder models.

In developing the methodology, we hypothesized that there is a dynamical low-dimensional manifold – state-process – present in the data, where its estimation will help to better understand the relationship between complex and distributed brain dynamics and behavioral readout. In our solution, we assumed that the dimension of the state is pre-known; however, identifying the dimension of the state process is of great interest. There are fewer established solutions that provide principle solutions to find the dimension of the state. In the auto-encoder model, we set the dimension of the state as a part of the behavioral observation process model. However, it is still of great importance to assess the optimal dimension of the state process. Note that we derive the posterior distribution of the state given the data, and we can study attributes of the posterior distribution to increase/decrease dimension of the state. Another possible solution is defining multiple state processes with independent noise processes; we also set a sparse prior on these states’ noise processes and use the Bayesian posterior to find posteiro of their noise variance. The noise variance posterior will help in search for optimal dimension. This will be one of our future research directions.

For D-D direct and auto-encoder model, we focused on deriving model identification and state estimation processes. We used BIC and other model selection algorithms to pick a proper history term for the observation processes. However, we did not discuss the goodness-of-fit process to better examine the extent of the model fit to data. Utilizing simulation data and using goodness-of-fit techniques to better assess the model fit is another research direction we pursue to better address the pros and cons of proposed framework.

Here, we mainly focused on the model development and how the proposed techniques will help to analyze complex and high dimensional data like the conversation dataset. Understandingthe neural mechanisms of those cognitive processes has significant prioritiess for experimental and clinincal neuroscientist, which is not discussed in this research. We puruse another research to better understand underlying neural mechanisms of intention using the inference being drawn from our modeling results.

## Acknowledgments

We thank Prof. Loren Frank and Dr. Anna K. Gillespie from the University of California at San Francisco who provided spiking activity dataset for decoding movement trajectory problem (sections 3.A and 3.B).

## Appendix A. Moving to a Fully B-DNN

Let’s assume that there is a prior probability over the network parameters **Ω**, *p*(**Ω**). Now, we require to build the joint posterior over the states value and the DNN model parameters defined by *p*(***x***_0:*k*_, **Ω**|***s***_1:*k*_, ***z***_1:*k*_). The posterior distribution is proportionate to full likelihood of the observation which can be defined by

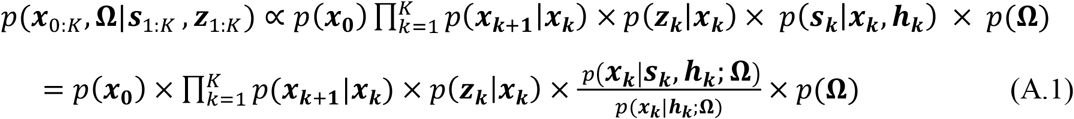

where ***z***_***k***_ and ***s***_***k***_ are the neural activity and the behavioral observations at the time index *k*; respectively. Note that constant term will be *p*(***s***_1:*k*_, ***z***_1:*k*_); as we see later, this term is not needed in our DNN training process.

Given the fact that the exact solution for *p*(***x***_0:*k*_, **Ω**|***s***_1:*k*_, ***z***_1:*k*_) due to a large dimension of **Ω** is intractable, we can approximate the true posterior *p*(***x***_0:*k*_, **Ω**|***s***_1:*k*_, ***z***_1:*k*_) by a variational distribution defined by

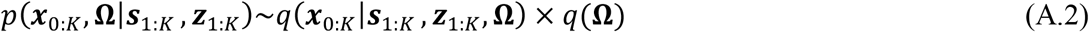

Where *q*(**Ω**) has a known functional form with a set of fixed parameters. Note that, we have already derived the solution for the first term on the right side of equation (A.2), which is the smoother solution with a known model parameter set. Equation (7) provides the smoother solution with neural observation; this solution can be extended for observation process including both neural and behavioral data. To find the proper set of parameters for the DNN weights’ distribution, we can minimize the Kullback-Leibler (KL) divergence between *q*(***x***_0:*k*_|***s***_1:*k*_, ***z***_1:*k*_, **Ω**) × *q*(**Ω**) and the true posterior *p*(***x***_0:*k*_, **Ω**|***s***_1:*k*_, ***z***_1:*k*_) [51]. The optimization corresponds to minimizing the following metrics

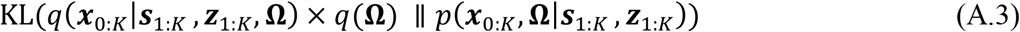

where, we require to find new set of paramaters for a specific functional form of *q*(**Ω**). As we discussed *q*(***x***_0:*k*_|***s***_1:*k*_, ***z***_1:*k*_, **Ω**) can be replaced by *p*(***x***_0:*k*_|***s***_1:*k*_, ***z***_1:*k*_, **Ω**), which we already have a solution for it.

The KL described in equation (A.2) can be written as

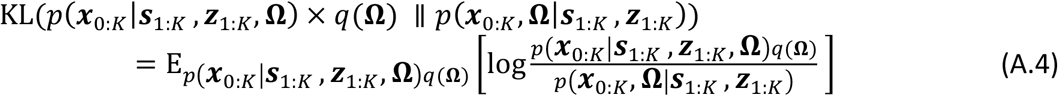

We can rewrite the KL distance as – moving forward, for simplicity, we use KL without its arguments.

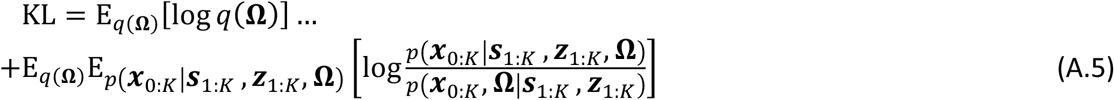

We can replace *p*(***x***_0:*k*_,**Ω**|***s***_1:*k*_, ***z***_1:*k*_) term using equation (A.1).

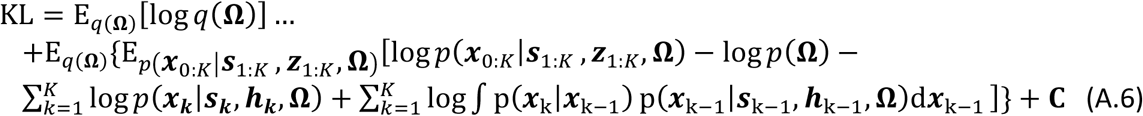

where, the terms which are not a function of the DNN weights’ parameters are represented by C. Let’s assume that **Ω** distribution paramaters are defined by ***ω*** and ***ω***^(**0**)^ is our initial guess of these parameters defining *p*(**Ω**; ***ω***^(**0**)^). Our objective is to minimize KL by tuning ***ω*** parameters along with other paramaters of the auto-encoder model. We optimize other paramaters of the model using the iterative EM algoirthm. As a result, KL needs to be minimized iteratively as the model paramaters and state estimation are updated per eahc iteration of EM step. With this assumption, we can rewrite the KL by

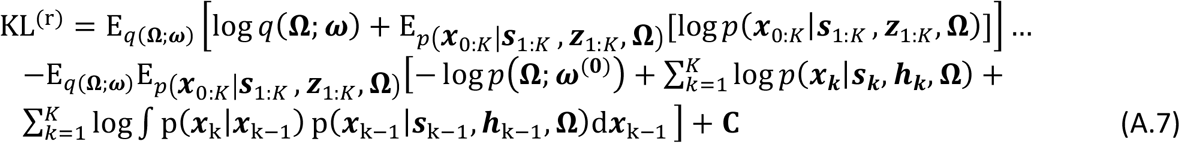

To minimize KL^(r)^, we require to find a new set of paramaters which are called, ***ω***^(*r*)^. Solving this optimization problem is computationally interactable; as a result, we provide a suboptimal solution using the EM solution we already proposed to find ML estimate of the DNN weights. As we discussed, this solution might be a proper apporach when the numer of weights in DNN is limited or the DNN weights are correlated. The solution is similar to a coordinate descent optimization, where we assume the auto-encoder models are updated recursively. Some of recent efforts to address Bayesian learning in dynamical auto-encoder model can be found in [40].

Let’s assume we have *q*(**Ω**; ***ω***^(r)^) for the *r*^t*h*^ iteration; we can define

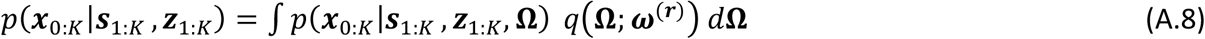

Let’s assume, we draw *D* samples from *q*(**Ω**; ***ω***^(*r*)^), *d* = 1 … *D*; using these samples, we can find the posterior estimation of state given behavioral and neural data. Note that, 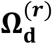 is *d*^t*h*^ sample of DNN network weights at iteration *r*. In other word, we have *D* DNN networks, where for each network, we can have 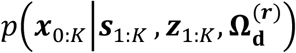 and the state posterior is defined as weighted sum of these distribution

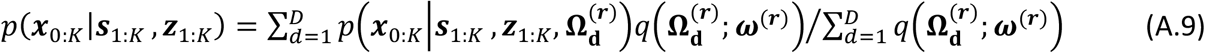

To calculate (A.9), we require to have *q*(**Ω**; ***ω***^(*r*)^).

To find *q*(**Ω**; ***ω***^(*r*)^), we assume there are *D* dynamical auto-ecnoder models and each model is trained using the EM alogithm defined in section **2.D**. Note that each DNN is trained using a regularizaion term defined by *p*(**Ω**; ***ω***^(**0**)^) – adding *p*(**Ω**; ***ω***^(**0**)^) to EM algorithm is straight forward. It is worth to mention that DNN training at iteration *r* can start with weights estimated at iteration *r* − 1. Training *D* DNN will provide new set of paramaters 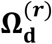 for each iteration; we can find updated paramaters for ***ω***^(*r*)^ through *D* DNN models, defined by

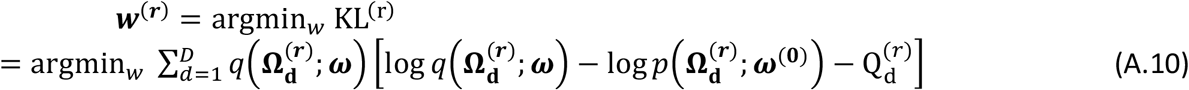

where, 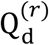 is defined by equation (22).

As a result, our B-DNN training includes *D* auto-encoder model trained in parallel. Using (A.10), we can find approximate posterior for DNN weights, and use that posterior to build a more robust posterior distribution of the state. When we use full B-DNN as part of our EM algorithm, we can replace the posteiror distribution in EM with the postrior distribution defined in equation (A.9), and use this distribution to find the state process and behavioral model paramaters.

